# Rethinking yield stability through phenotypic plasticity and its link to modern statistical methods

**DOI:** 10.64898/2026.01.07.698075

**Authors:** Victor Sadras, Matthew Welsh, Beata Sznajder, Julie Hayes, Matthew Reynolds, Julian Taylor

## Abstract

In the context of advocacy for yield stability, trade-offs between yield and yield stability, the frequent lack of definitions, and the variation in methods when they are explicit, we connect two perspectives: phenotypic plasticity and factor analytics. Phenotypic plasticity brings over a century of research in developmental biology, ecology and evolution, and is gaining traction in crop science. Factor analytics is an advanced linear mixed model of multi-environment data with factor analytic variance structures for the variety-by-environment interaction effect. We (1) review phenotypic plasticity, define *agronomically adaptive plasticity* where varieties (or practices) consistently return superior yield (or other traits) across environments with no trade-off, and describe percentile-plasticity plots to assess the agronomic value of plasticity; (2) outline factor analytic models, and (3) link plasticity and factor analytic models mathematically and empirically. We show that phenotypic plasticity of cereal yield correlates positively with overall performance obtained from factor analytic models when plasticity is adaptive and negatively when it is maladaptive. We conclude that phenotypic plasticity contributes biological meaning to opaque analytical approaches, and that biologically grounded statistics are needed to challenge weak agronomic narratives, such as advocacy for stability that might reflect decision biases rather than critical consideration of its benefits.

**Highlight:** Uncritical advocacy for crop yield stability is common. Here we advance a biological–statistical synthesis of yield stability and test theoretical predictions with actual wheat and oat yield data.

## Introduction

‘Balance’ and ‘stability’ are culturally embedded, value-laden concepts. In both Western cultures that draw deeply from ancient Greek philosophy and their Eastern counterparts informed by Taoism, Confucianism, and Buddhism, ‘balance’ and ‘stability’ are regarded as desirable (Head, 2025; Hwang, 2012; Mou, 2023; Mou *et al*., 2012; Simberloff, 2014; Yulie, 2015). Making explicit the cultural dimension of the concept, Head (2025) challenges the ‘balance-is-good’ proposition for Murray-Darling water allocation in the context of climate change. Russel Wallace, *ca.* 1855, challenged the notion of a balance of nature as an undefined entity whose accuracy could not be tested, leading to the contemporary rejection of the concept and a focus on a dynamic, often chaotic nature buffeted by constant disturbances (Simberloff, 2014). In ecology, the notion of balance in nature is recognised as a *panchreston*: a term that means so many different things to different people that it has become useless (Simberloff, 2014). In evolutionary biology, a paradox of terminology is apparent whereby ‘a language of stasis and stability has long dominated the science of change’ (p. 8, West-Eberhard, 2003). Stabilising selection, *i.e.,* selection against extreme values of a trait, is assumed to be common (p. 234, Odling-Smee, 2024) despite weak empirical evidence (Haller and Hendry, 2014). In decision making, a preference for stability is reflected in people’s widespread bias towards certainty (Box 1).

In agriculture, yield stability is a growing focus of research (Fig. 1A), and ‘balance’ and ‘stability’ are regularly invoked as desirable goals. As a reflection of the ‘stability-is-good’ notion, advocacy for yield stability with no definition or quantification is not rare (De Jonge *et al*., 2025; Derbyshire *et al*., 2024; Jian *et al*., 2024; Ravi *et al*., 2024; Stadnik *et al*., 2024; Wakjira *et al*., 2024; Xue *et al*., 2024). Lack of definitions is a source of ambiguity.

**Figure 1.**
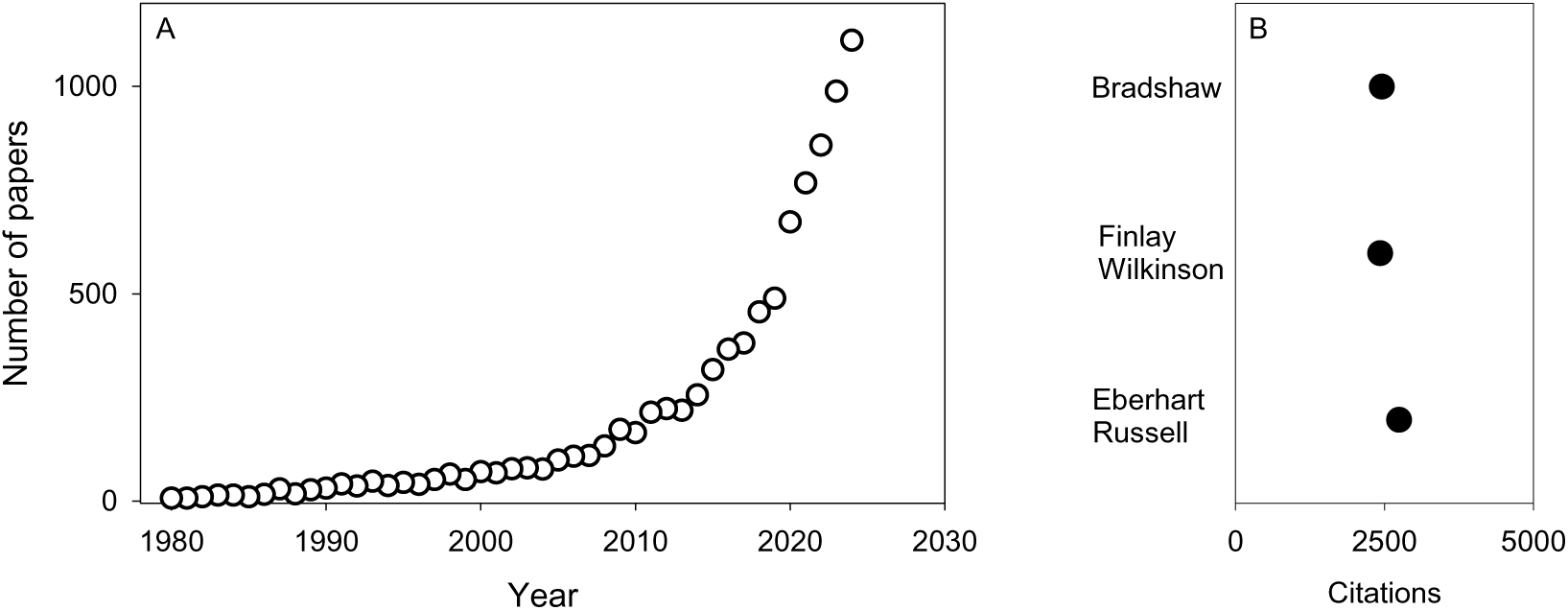
Yield stability in the scientific literature. (A) Number of papers retrieved from Web of Science searching for “yield” and “stability” and “crop” in all bibliographical fields, period 1980-2024. (B) Citations from Web of Science (August 2025) of influential 1960s papers dealing with relationships between genotype and environment in developmental and evolutionary (Bradshaw, 1965) and plant breeding contexts (Eberhart and Russell, 1966; Finlay and Wilkinson, 1963).

Subhavyuktha *et al*. (2024) reviewed parametric and non-parametric methods to calculate trait stability; hence, our aim is not to review methods but to illustrate the context and diversity of approaches. Searching Web of Science with key words ‘crop’, ‘yield’ and ‘stability’ in any field returned 1117 papers published in 2024 (August 2025). Some of them dealt with structural stability of cell membranes, photosynthesis, biochar, soil aggregates, depth of tillage, or atmospheric process (Albasha *et al*., 2024; Alves *et al*., 2024; Dwivedi *et al*., 2024; Holatko *et al*., 2024; Ramesh and Raghavan, 2024; Ye *et al*., 2024). Excluding these papers, and those where yield stability is not defined, we sampled papers where stability of yield and other agronomic traits is operationally defined*, i.e.,* the calculation method is explicit (Table 1). These studies ask how the stability of yield or other traits varies with genotype in a breeding context, and with agronomic practices such as fertilisation and cultivar mix; some studies look at yield stability with genotype and management intersecting with climate trends; identifying high yield, high stability genotypes and practices is a common goal. Agronomic studies use metrics such as coefficient of variation and *ad-hoc* indices. Additive Main Effects × Multiplicative Interaction (AAMI) models, slopes of reaction norms relating phenotype and environment, factor analytic models, and other methods to account for genotype-by-environment interactions are more typical of studies aimed at ranking phenotypes in pre-breeding and breeding (Table 1).

**Table 1.** Sample of papers retrieved searching Web of Science (August 2025) with key words ‘yield’ and ‘crop’ and ‘stability’ in all fields of the bibliographical record. Of the 1117 papers retrieved in 2024, we discarded papers that (i) dealt with structural stability of cell membranes, soil aggregates etc., and (ii) did not quantify stability. After this filtering, we list papers that represent the diversity in methods to calculate stability in agronomic (A) and breeding (B) contexts. No attempt was made to quantify the frequency of use of the methods or their reliability, but they are summarised here to illustrate their diversity.

| Context | Method to quantify stability | Focus | Source |
| --- | --- | --- | --- |
| A | standard deviation, SD | relationship between volatility of combined yields and diversity for standardised yield data of major crops grown in Germany between 1977 and 2018 | (Ahrends *et al.*, 2024) |
| A | coefficient of variation, CV | effects of biochar application on crop yield variability | (Zhang *et al.*, 2024b) |
| A | coefficient of variation, CV | effect of shallow groundwater on US maize yield and its stability, 1999-2018 | (Deines *et al.*, 2024) |
| A | coefficient of variation, CV | effect of insect pollination on yield, quality and trait stability of oil tree peony | (Zhang *et al.*, 2024a) |
| A | inverse of the coefficient of variation | yield and yield stability of oat-soybean strip intercropping | (Feng *et al.*, 2024b) |
| A | inverse of the coefficient of variation | yield stability of perennial and annual cropping systems | (Shang *et al.*, 2024) |
| A | absolute stability ratio:<br>$\ln SD_t - SD_c$<br>relative stability ratio:<br>$\ln CV_t - \ln CV_c$<br>t is treatment, c is control | Meta-analysis of biomass and allocation comparing a warming, warming plus CO<sub>2</sub> fertilisation, and warming plus organic amendment | (Wang *et al.*, 2024b) |
| A | standard deviation,<br>first-order lower partial moments of a yield anomaly time series | long-term variation in yield and yield stability of wheat and maize in response to climate and NPK supply | (Zhu *et al.*, 2024) |
| A | coefficient of variation,<br>POLAR (POwer LAw Residuals) stability index, which removes the dependence of variance on mean yield | yield and yield stability of cultivar mixtures | (Huang *et al.*, 2024) |
| A | sustainable yield index<br>$SYI = (Mean\ Yield - Yield\ SD) / Y_{max}$ | crop yield and its stability under prolonged application of chemical fertilisers and/or manure | (Wang *et al.*, 2024a) |
| A | coefficient of variation,<br>sustainable yield index | organic substitution ratio in vegetable production | (Xu *et al.*, 2024) |
| A | coefficient of variation,<br>sustainable yield index | yield and yield stability of cotton in response to nitrogen fertilisation | (Feng *et al.*, 2024a) |
| A | coefficient of variation,<br>sustainable yield index | wheat yield in response to rotation and nitrogen fertilisation | (Yan *et al.*, 2024) |
| A | coefficient of variation,<br>sustainable yield index | yield and yield stability of rapeseed after maize and maize-legume intercrop | (Yang *et al.*, 2024) |
| B | GGE biplot | yield and yield stability of potato varieties | (Ahmed *et al.*, 2024a) |
| B | additive main effects and multiplicative interaction (AMMI) model | high yield, high yield stability potato cultivars | (Tatarowska *et al.*, 2024) |
| B | AMMI | high yield, high yield stability maize hybrids | (Haripriya and Radhika, 2024) |
| B | Bayesian AMMI model | genotype-by-environment patterns of chickpea yield and days to emergence | (Din *et al.*, 2024) |
| B | AMMI stability value (ASV), quantifying the genotype's interaction with environment, and genotype stability index, which incorporates both stability and mean performance across environments | genotype-by-environment interaction and cultivar stability in vegetable soybean (edamame) | (van der Merwe *et al.*, 2024) |
| B | AMMI Stability Value and stability index<br>Stability variance (Shukla, 1972)<br>Eberhart and Russell (1966) slope of reaction norm | high yield, high yield stability rice | (Ahmed *et al.*, 2024b) |
| B | multi-trait stability index, MTSI | high erucic acid *B. carinata* genotypes with stable performance in multiple desirable traits | (Tesfaye *et al.*, 2024) |
| B | Eberhart and Russell (1966): slope of reaction norm and mean square deviation from regression $S^2d_i$ | stability for nitrogen fixation and seed yield in mungbean genotypes | (Arya *et al.*, 2024) |
| B | Eberhart and Russell (1966): slope of reaction norm and mean square deviation from regression $S^2d_i$<br>AMMI stability value, ASV | high yield, high yield stability in faba bean | (Wondaferew *et al.*, 2024) |
| B | Eberhart and Russell (1966): slope of reaction norm and mean square deviation from regression $S^2d_i$<br>Stability value (ASV) and yield stability index (YSI) from AMMI | high yield, high yield stability rice | (Roy *et al.*, 2024) |
| B | Eberhart and Russell (1966): slope of reaction norm and mean square deviation from regression $S^2d_i$<br>AMMI stability parameter ASTAB, stability index ASI, stability value ASV | high yield, high yield stability rice | (Abd El-Aty *et al.*, 2024) |
| B | AMMI stability value ASV and genotype stability index | genotype-by-environment interaction and cultivar stability in vegetable soybean (edamame) | (van der Merwe *et al.*, 2024) |
| B | Stress susceptibility index | QTL for wheat yield stability under drought | (Liu *et al.*, 2024) |
| B | Factor analytic overall performance and stability metrics | Analysis of three industry relevant groundnut traits | (Coulibaly *et al.*, 2024) |
| B | Factor analytic overall performance and stability metrics | Analysis of nematode resistance in wheat | (Rognoni *et al.*, 2024) |

In the context of advocacy for yield stability that might involve cultural and cognitive bias (Box 1), trade-offs between yield and stability of yield (Calderini and Slafer, 1999; Giordano *et al*., 2025b; Raj *et al*., 2023), the frequent lack of definitions, and the variation in methods when they are presented, this paper connects two perspectives: phenotypic plasticity and factor analytical models embedded in linear mixed models (Smith *et al*., 2001; Smith *et al*., 2005). Our aim is to highlight the value of combining biological thinking and pragmatic statistics. Phenotypic plasticity draws on more than a century of research in developmental biology, ecology and evolution, and is increasingly used to frame questions in crop science. Factor analytics is a contemporary method to analyse multi-environment data involving a linear mixed model with factor analytic variance structures for the variety-by-environment interaction effect. We review biological insights from phenotypic plasticity, advance a definition of agronomically adaptive plasticity, describe percentile-plasticity plots to probe for the adaptive value of plasticity, outline factor analytical approaches, and link phenotypic plasticity and factor analytic models mathematically and empirically. Using published oat and wheat yield data, we show that phenotypic plasticity calculated as the slope of reaction norms correlates positively with overall performance from factor analytics when plasticity is adaptive and negatively when it is maladaptive.

### Phenotypic plasticity

#### Overview

Phenotypic plasticity, *i.e.,* ‘the amount by which the expressions of individual characteristics of a phenotype are changed by different environments’ (Bradshaw, 1965) is central to developmental biology, ecology, and evolution (Day and McLeod, 2018; DeWitt and Scheiner, 2004; Diouf *et al*., 2020; Mitchell *et al*., 2024; Nicotra *et al*., 2010; West-Eberhard, 2003). Early in the 20^th^ century, Woltereck (1909) measured morphological traits in water flea clones grown under contrasting environmental conditions, and drew curves, which he called reaction norms, relating traits and environmental factors; Sarkar (2004) traced the evolution of the concept since its formulation. The slope of reaction norms quantifies phenotypic plasticity, which can also be calculated with other measures of variation including range, variance ratio, relative change between treatments, *e.g.,* drought vs well-watered, and random regression mixed models (Berlin *et al*., 2017; Bhatia *et al*., 2014; Dingemanse *et al*., 2010; Kadam *et al*., 2017; Ruiz *et al*., 2019; Sadras and Richards, 2014; Valladares *et al*., 2000){Arnold, 2019 #23224}. Some methods return similar rankings of phenotypic plasticity, *e.g.,* plasticity calculated as variance ratio correlates with slopes of reaction norms in some data sets (Sadras and Richards, 2014). Metrics based on differences or ratios return a single value of plasticity, whereas reaction norms can account for non-linearity (Woltereck, 1909). In controlled environments where a single source of environmental variation dominates, tight reaction norms can be fitted, *e.g.*, between leaf expansion rate and vapour pressure deficit (Reymond *et al*., 2003). In the field where multiple environmental factors often influence the phenotype, reaction norms are bound to be scattered (Cade and Noon, 2003) and can be tightened using the mean of the trait as the environmental variable (Calò *et al*., 1975; Curin *et al*., 2021; Trentacoste *et al*., 2011).

Phenotypic plasticity has been interpreted through the lens of adaptation to environmental heterogeneity, where four strategies have evolved (DeWitt and Langerhans, 2004): (1) specialisation, whereby a single phenotype is produced that is well adapted to a particular environment even though it may experience a range of environments; (2) generalisation, whereby a ‘general purpose’ phenotype is produced, with moderate fitness in most environments; (3) bet-hedging, whereby an organism produces either several phenotypes (*e.g.*, among units in a modular plant) or a single phenotype probabilistically; and (4) phenotypic plasticity, whereby environmental signals trigger the production of alternative phenotypes. Modelling these four strategies under the assumption of ‘perfect plasticity’ and a simplified range of environments returned a ratio of fitness after four generations of 1 : 1.6 : 1.5 : 25 (DeWitt and Langerhans, 2004). The conclusion is that in the absence of constraints, plasticity is always superior in variable environments. The lack of ubiquity of plasticity, however, highlights the existence of ubiquitous constraints including the cost of plasticity, developmental constraints, and the unreliability of environmental cues that guide development (Aphalo and Sadras, 2021; Auld *et al*., 2010; De Witt *et al*., 1998; Murren *et al*., 2015; Odling-Smee, 2024; Volis, 2009). Phenotypic plasticity can be therefore adaptive or maladaptive (see *Definition: agronomically adaptive plasticity)*.

Contemporary to Bradshaw’s (1965) review of phenotypic plasticity, Finlay and Wilkinson (1963) and Eberhart and Russell (1966) looked at genotype-dependent response to the environment in a crop breeding context; all three studies have been and remain influential (Fig. 1B). Eberhart and Russell (1966) fitted linear models relating the yield of a variety and an environmental index:

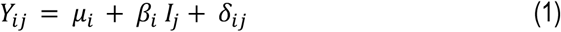

where *Y*_*ij*_ is the yield of variety *i* at environment *j*, *μ*_*i*_ is the mean yield for variety *i* across environments, *β*_*i*_ is the regression coefficient that measures the response of variety *i* to varying environments, *I*_*j*_ is the environmental index calculated as the mean of all varieties at environment *j* minus the grand mean, and *δ*_*ij*_ is the deviation from regression of variety *i* at environment *j*; the deviation *δ*_*ij*_ is squared and summed to return a stability parameter 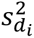. A desirable phenotype has been proposed that features high mean yield, *β*_*i*_ close to 1, and small 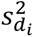 (Eberhart and Russell, 1966).

Finlay and Wilkinson (1963) were aware of research on phenotypic stability, heterosis, developmental homeostasis, and response to environmental change in developmental biology, population genetics and evolutionary studies, but were concerned with the relevance of laboratory studies for field crops. Using a log-log model for statistical reasons, they made explicit that ‘means of yields on a logarithmic scale correspond to geometric means on the natural scale’ (Finlay and Wilkinson, 1963); geometric means have been used as a measure of fitness in evolution (Levins, 1968; Wimsatt, 2006). Both Finlay and Wilkinson (1963) and Eberhart and Russell (1966) fitted reaction norms where the slope is phenotypic plasticity, but the connection was not made explicit at the time. It would take another four decades for crop science to embrace phenotypic plasticity as a useful angle to look at the genotype-dependent phenotypic response to the environment (Adams *et al*., 2025; Bicego *et al*., 2024; D’Andrea *et al*., 2013; de Felipe and Alvarez Prado, 2021; Elazab *et al*., 2025; Giordano *et al*., 2024; Govta *et al*., 2025; Iqbal *et al*., 2024; Kadam *et al*., 2017; Otwani *et al*., 2023; Peltonen-Sainio *et al*., 2011; Prohaska *et al*., 2024; Sadras and Lawson, 2011; Sadras *et al*., 2009; Sadras and Trentacoste, 2011; Schell *et al*., 2025; Zhao *et al*., 2025). With a focus on plant breeding for drought adaptation, Cooper and Messina (2022) illustrate a shift in mindset from the relationship between genotype and environment seen as a nuisance to one where it is a ‘subject of productive interest’ (West-Eberhard, 2003).

The early prediction that ‘the plasticity of a character is an independent property of that character and is under its own specific genetic control’ (Bradshaw, 1965) has been demonstrated experimentally (Alvarez Prado *et al*., 2014; Berlin *et al*., 2017; Bloomer *et al*., 2014; Giordano *et al*., 2025a; Kadam *et al*., 2017; Marguerit *et al*., 2012; Reymond *et al*., 2003). This has three practical implications. First, probing for the genetics of trait response to environment pre-empts QTL-by-environment interactions (Kadam *et al*., 2017; Reymond *et al*., 2003). Second, phenotypic plasticity and the trait *per se* can be partially independent targets of breeding and selection. Third, if stability is desirable and plasticity is its reverse, breeding and selection for agronomic adaptation should favour more stable, less plastic phenotypes; but selection for yield and agronomic adaptation has favoured high phenotypic plasticity of yield in many breeding programs: wheat in Australia, US, Argentina, UK and Canada (Giordano *et al*., 2025a), oat in Australia (Hayes *et al*., 2026), soybean in Argentina (de Felipe and Alvarez Prado, 2021), and spring wheat in Finland (Peltonen-Sainio *et al*., 2011). The apparent contradiction between high plasticity and stability reinforces the need for unambiguous definitions and rigorous biological interpretation of statistical models.

The relative rate of increase of average yield of maize, rice and wheat is slowing down in major production countries, with consequences for food security (Grassini *et al*., 2013). In this context, Sadras (2024) has linked developmental (Amzallag, 2000) and ecological and evolutionary (Berg, 1960; Terentjev, 1931) perspectives on phenotypic integration leading to the prediction that breeding favours high plasticity of yield, where it is adaptive (see definition below), at the expense of heritability. Experimental evidence for this prediction has been reported for winter wheat in the US, where broad-sense heritability of yield has declined from 0.23 to 0.15 between 1990 and 2020 in parallel to a linear increase in phenotypic plasticity of yield (Giordano *et al*., 2025a). Phenotypic plasticity linked with phenotypic integration thus identified reduced heritability as an overlooked factor contributing to the reported slowing down of genetic gain in yield.

#### Agronomically adaptive plasticity

We need a definition to conclude objectively whether phenotypic plasticity is agronomically desirable. Here we identify *agronomically adaptive plasticity* (AAP) where genotypes (or practices) consistently return superior yield (or other traits) across environments with no trade-off. Our definition resonates with Maynard Smith’s (1982) definition of an evolutionary stable strategy (ESS) for the evolutionary game as a strategy ‘such that, if all members of a population adopt it, no mutant strategy can invade.’ Both ESS and AAP are unrealistic targets, *i.e.*, phenotypes cannot be optimised in nature (Darwin, 1859) or agriculture (Sadras and Denison, 2016), but are useful null hypotheses as departures from ESS and AAP predictions would reveal adaptive constraints (DeWitt and Langerhans, 2004; Gómez-Llano *et al*., 2025; Kinraide and Denison, 2003; Sadras and Denison, 2016). This parallels Tversky and Kahneman’s (1974) ‘heuristics and biases’ perspective in decision making (Box 1).

#### Percentile-plasticity plots to test agronomically adaptive plasticity

Percentile-plasticity plots are useful to visualise and assess the agronomic value of phenotypic plasticity (Fig. 2). For each genotype, trait percentiles, say 90^th^ and 10^th^, are calculated to quantify the trait, for example yield, under favourable and unfavourable (*e.g.*, severe drought), conditions; percentiles account for contrasting conditions while reducing the impact of outliers. Percentiles plotted against plasticity, calculated with any of the methods above, return patterns as illustrated in Fig. 2, which can be tested against our definition of agronomically adaptive plasticity. In pattern A, the more plastic pink phenotype is superior in good conditions, with no trade-off under stress; high phenotypic plasticity is agronomically adaptive. Pattern B is the opposite: plasticity is agronomically maladaptive because the blue phenotype, with the lowest plasticity, is superior under stress with no trade-off under favourable conditions. Pattern C involves a trade-off where the pink phenotype is superior under favourable conditions at the expense of stress adaptation. Selection for the trait of interest should favour high phenotypic plasticity in A and low plasticity in B. In C, additional criteria are needed to solve the trade-off.

**Figure 2.**
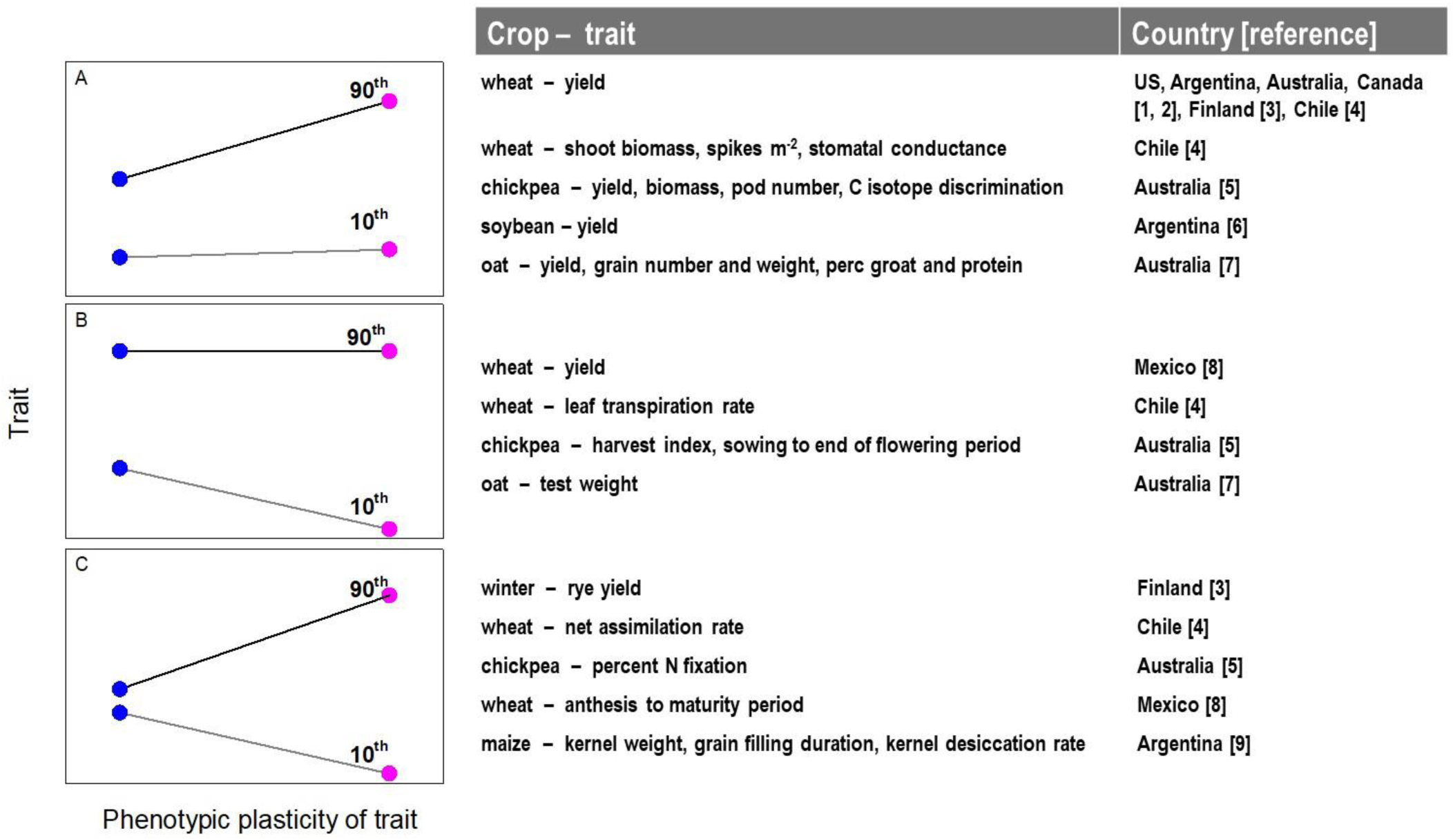
Testing the agronomic value of phenotypic plasticity. Percentile-plasticity plots highlighting two extreme phenotypes: blue, with the lowest phenotypic plasticity, and pink, with the highest phenotypic plasticity. A sample of three patterns is shown where (A) the pink phenotype is superior under favourable conditions (90^th^ percentile, black line) with no trade-off under stress (10^th^ percentile, grey line); (B) the blue phenotype is superior under stress with no trade-off under favourable conditions; (C) the pink phenotype is superior under favourable conditions and inferior under stress. Experimental evidence supporting the patterns in A-C: [1] Giordano et al. (2024), [2] Giordano et al. (2025a), [3] Peltonen-Sainio et al. (2011), [4] Elazab et al. (2025), [5] Sadras et al. (2016), [6] de Felipe and Alvarez Prado (2021), [7] Hayes et al. (2026), [8] Sadras et al. (2009), [9] Alvarez Prado et al. (2014).

### Factor analytic models

#### Outline

Factor analytic (FA) models are the random-effects analogue of the Additive Main Effects × Multiplicative Interaction (AMMI) model commonly used in plant breeding (Table 1, and Malosetti *et al*., 2013). An FA model is a term used to explain a general linear mixed-model (LMM) that has an embedded factor analytic model for the genotype-by-environment (GxE) interaction. In the following sections, we discuss the FA model for the GxE interaction in a statistical context; in the Discussion, we explain why GxE is a biologically inappropriate description of development. We base our outline on Smith and Cullis (2018) and others (Chaves *et al*., 2024; Fradgley *et al*., 2025) that use a LMM of the form

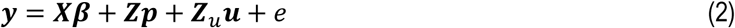

where ***y*** is the raw trait data, ***β*** is a set fixed effects that consists of effects to estimate experiment means, ***p*** are a set of non-genetic random effects usually linked to environmental variation and *e* are structured model residuals. In this model we focus on 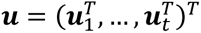 as a set of genetic random effects for *m* genotypes in *t* environments that are decomposed into a factor analytic model of order *k*. For the random effect of individual *i* in environment *j*, the model can be expressed as

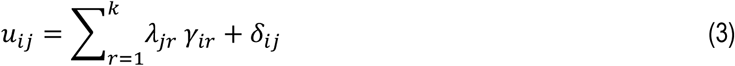

where *λ*_*jr*_ is the loading of environment *j* on factor *r*, *γ*_*ir*_ is the score for the *i*th genotype on factor *r*, and *δ*_*ij*_ are model residuals that are assumed to have a distribution 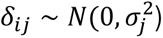 where 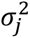 is a specific variance associated with the *j*th environment. Note that as experiment means are estimated through *β*, the random effects *u*_*j*_, ∀ *j* = 1,…, *t* are zero centred.

In vector-matrix form, eq. 3 can be specified as

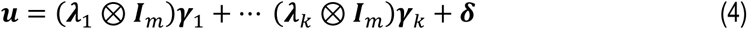

where, ***λ***_*r*_ and ***γ***_*r*_ are the loadings and score vectors for the *r*th factor and 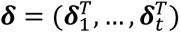 is the residual vector. This allows the variance-covariance of the genetic effects to be immediately specified as

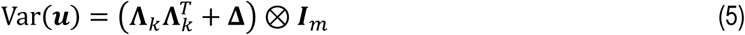

where **Λ**_*k*_ = [***λ***_1_… ***λ***_*k*_] is the *m* × *k* matrix of loadings and Δ is a diagonal matrix with (*ij*)th diagonal element 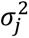. The complete set of loadings in **Λ**_*k*_ are unobserved or latent covariates that require estimating, alongside the variance parameters in **Δ**. Structurally, equation (4) indicates that 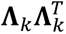 forms the GxE covariance of the genetic effects common across the environments whereas **Δ** represents the site-specific variation that cannot be accounted for by the common GxE component.

In general, sophisticated LMM solvers are required to appropriately estimate the variance parameters embedded in **Λ**_*k*_, **Δ**. A well-established approach is to use Residual Maximum Likelihood (REML) of Patterson and Thompson (1971) available in linear mixed modelling software ASReml-R (Butler *et al*., 2018). After fitting, loadings in Λ_*k*_ are typically rotated to a principal-component form so that factor 1 explains most of the covariance with factor 2 orthogonal to factor 1 and other factors following similarly.

#### Interpretation

Empirically evidence for positive FA1 loadings after rotation of loadings suggests they are linked to performance of the trait across environments,, leading to the development of Factor Analytic Selection Tools, FAST (Smith and Cullis, 2018). Consider an alternative form for the random effect of individual *i* in environment *j*

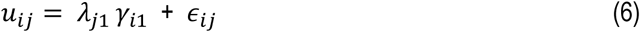

where ***∊***_*ij*_ becomes a new residual that subsumes the factor analytic components for the factors *k* > 1 and *δ*_*ij*_. Then, empirical measures of overall performance (OP) and stability (ST) for each of the genotype across all environments from the FA1 are defined for individual *i* (Smith and Cullis, 2018):

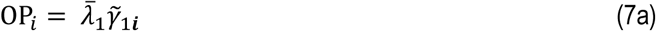

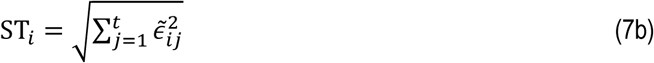

where 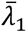 is the average of all the positive loadings of the first factor. The OP for each genotype can then be considered as the fitted value of the genotype regression at the average value of the latent covariate, and ST is a measure of the spread of the residuals from this regression. These two measures provide a complete interpretability of genotype performance. Positive OP indicates better than average performance of a genotype across environments and negative OP indicates below average performance. As ST measures spread from the genotype regression, low ST suggests stability across environments. Both OP (eq. 7a) and ST (eq. 7b) are on the same scale so they can be combined to rank each genotype with an index I_i_, for example

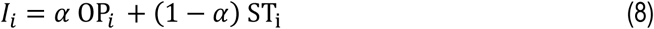

This equation makes explicit a trade-off between overall performance and stability, with ***α*** = 1, solely selecting for performance and, ***α*** = 0, solely selecting for stability. Chaves *et al*. (2024) applied a FAST-based index, including a reliability weight, to select *Eucalyptus* clones with both high OP and high stability across environments; they went beyond a nominal classification of environments (location-year) to identify ambient temperature, radiation, and soil characteristics as key environmental factors associated with relevant traits.

### Linking phenotypic plasticity and factor analytic models

#### Mathematically

Phenotypic plasticity and the factor analytic model can be linked mathematically with minimal assumptions. Consider an alternative form for (1) where we assume that an observed environmental index 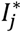 is such that when its equal to zero the trait value tends to zero also. An example of this index could be the environmental means for the trait being analysed. With these assumptions the main effect of the genotypes *μ*_*i*_ → 0, ∀ *i* = 1,…, *m*. Note, this has no material change on *β*_*i*_, ∀ *i* = 1,…, *m*. Now let 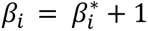 then eq (1) can be rewritten as

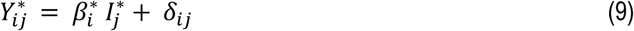

where 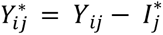. This is equivalent to sweeping out the trial means. This new regression mimics the form and function of the factor analytic model in eq. (6) for individual *i* in environment *j* that was used to generate the OP and ST of the genotypes. Thus, if the first-factor loadings in eq. (6), *λ*_*j*1_, ∀ *j* = 1,…, *t* align to scale with the environmental index 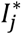, ∀ *j* = 1,…, *t* in (7), then the genotype scores *γ*_*i*1_, ∀ *i* = 1,…, *m* will be similar to the slopes 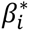, ∀ *i* = 1,…, *m* and, by default, *β*_*i*_, ∀ *i* = 1,…, *m*. This link is the reason for the FA1 component of the FA model to be seen as the principal ‘reaction norm’ axis in the absence of known covariates. This has been noted in detail by Tolhurst *et al*. (2022), who illustrate that the first latent factor captures only non-crossover (scale) GxE, while higher factors in the FA model capture crossover GxE.

From (7a), OP for the genotypes is calculated by scaling the empirical FA1 scores, 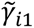, ∀ *i* = 1,…, *m* by the average of the estimated rotated FA1 loadings. This means that if the estimated FA1 loadings mimic the environmental index then the complete set of genotype overall performance OP_*i*_ ∀ *i* = 1,…, *m* is likely to be correlated with plasticity empirically calculated from 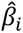, ∀ *i* = 1,…, *m*. This link is tested empirically in the next section.

#### Empirically

We tested the links between phenotypic plasticity and factor analytics outputs using published yield data for Australian oat comprising 128 cultivars and advanced lines grown in 106 environments (Hayes *et al*., 2026) and for 169 CIMMYT wheat lines grown in six contrasting, irrigated and rainfed environments in Mexico (Sadras *et al*., 2009). Yield ranges were similarly broad in both data sets: from 0.2 to 7.8 t ha^-1^ in oat, and from 0.4 to 8.1 t ha^-1^ in wheat. These data sets were selected to illustrate agronomically adaptive (Fig. 3A) and maladaptive (Fig. 3 B) plasticity of yield. Phenotypic plasticity was calculated as the slope of linear reaction norms and factor analytical models using the approach above were used to calculate overall performance and stability (eq. 7ab). A percentile-plasticity plot showed yield of oat phenotypes with high yield plasticity was markedly superior under favourable conditions and slightly better under stress (Fig. 3A). High phenotypic plasticity was neutral for wheat yield under favourable conditions and was associated with low yield under stress in our chosen dataset (Fig. 3B). Where phenotypic plasticity of yield was agronomically adaptive, *i.e.,* for oat in Australia, phenotypic plasticity correlated positively with overall performance from factor analytics (Fig. 3E). Where phenotypic plasticity was maladaptive, *i.e.,* for wheat in Mexico, plasticity correlated negatively with overall performance from factor analytics (Fig. 3D). The standard error of slopes from fitted reaction norms correlated with yield stability calculated from factor analytics, more strongly in the oat (r = 0.59, p < 0.0001) than in the wheat data set (r = 0.21, p = 0.005).

**Figure 3.**
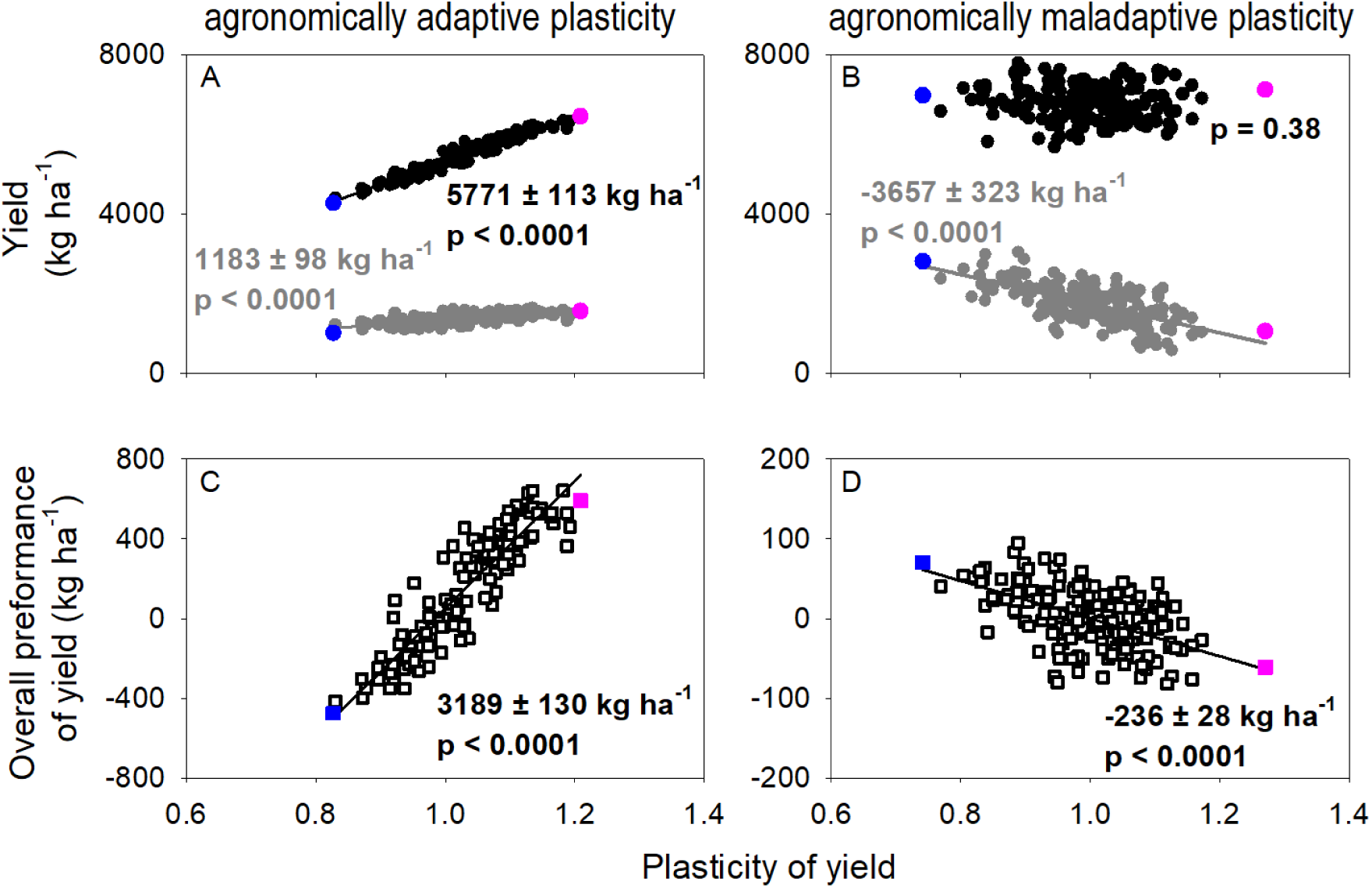
Linking phenotypic plasticity and factor analytics empirically. Percentile-plasticity plots show (A) phenotypic plasticity of grain yield was agronomically adaptive in a set of 128 oat cultivars and advanced lines grown in 106 Australian environments, and (B) maladaptive for yield of 169 CIMMYT wheat lines grown in six rainfed and irrigated Mexican environments. Black points are 90^th^ percentile, grey points are 10^th^ percentile. Phenotypes with the highest (pink) and the lowest (blue) phenotypic plasticity for yield are highlighted. Genotype-dependent phenotypic plasticity of yield was calculated as the slope of linear reaction norms. Overall performance from factor analytics (eq. 7a) correlated positively with (C) phenotypic plasticity where plasticity was adaptive and (D) negatively where it was maladaptive. In (A-D), numbers are slope ± standard error of lineal regressions (solid lines). Calculated from oat data in Hayes et al. (2026) and wheat data in Sadras et al. (2009).

## Discussion

A long list of scientific papers uncritically advocating for undefined yield stability is consistent with our proposition that preferences for balance and stability partially reflect cognitive tendencies and cultural influences (Introduction, Box 1). Demonstration of cognitive bias in the advocacy for yield stability in agriculture is beyond the scope of our paper, but awareness of how culturally embedded tendencies can skew our thinking is the first step towards acknowledging, accounting for, and potentially mitigating these biases.

The widely used Genotype x Environment x Management (GxExM) framework to interpret plant phenotypes in agricultural contexts has three limitations that can be partially ascribed to cognitive bias. Firstly, the genotype-by-environment interaction is a justified component of the phenotypic variance in statistical models but is a biologically misleading description of development because ‘genes do not interact directly with the external environment during development. All interaction is indirect, via effects of both factors in a preexisting phenotype…’(p. 15, West-Eberhard, 2003). Biologically, GxE is better described as genotype-dependent phenotypic response to the environment; this is a grammatical mouthful but illuminates a misconstructed concept. Secondly, the framework where the plant phenotype is seen as a passive consequence of genetic, environmental, and management factors overlooks that the phenotype modulates the genome, the environment in a process of niche construction, and indirectly influences management through the interplay with the phenotype of the farmer (Sadras and Hayman, 2025); a cautious teleonomic view of the phenotype is useful (Corning *et al*., 2023; Odling-Smee, 2024; Schrödinger, 1958). Third, the term ‘interaction’ in the GxExM framework introduces a bias against additivity, *i.e.,* lack of interaction (Côté *et al*., 2016; Jimenez *et al*., 2024; Ploschuk *et al*., 2025; Sadras *et al*., 2024; Sadras *et al*., 2025). Scientific publications that claim synergies with no empirical evidence are twice as likely than unsubstantiated claims of additivity and antagonism, and this bias has been ascribed to the more compelling narrative offered by synergies; synergies are more likely to make popular news headlines and papers referring to synergies return higher rates of citations than those reporting additivity and antagonism, especially for review articles (Côté *et al*., 2016).

Issues in the field of yield stability in agriculture include the lack of definitions, trade-offs between yield and yield stability, sophisticated statistics with opaque biological meaning, and weak agronomic narratives such as the advocacy for stability that might reflect decision biases rather than critical consideration of its benefits. Trade-offs between yield and yield stability are important. A factorial experiment that compared the yield of 20-55 cultivars under standard grower practice and intensive management in 21 environments in the southern US Great Plains demonstrated an agronomic trade-off where high yield stability was achieved at the expense of a high yield gap (Giordano *et al*., 2025b). In a breeding context, eq. (8) makes explicit this trade-off captured in α between 1, solely selecting for performance to 0, solely selecting for stability.

We advance a definition of agronomic adaptive plasticity and use percentile-plasticity plots to show both adaptive and maladaptive plasticity are trait-dependent and may emerge depending on the combination of genotypes and environments sampled (Fig. 2). To reach a conclusion of practical value, for instance to inform breeding decisions, large samples are needed that capture the diversity of genotypes and environments of interest (Hayes *et al*., 2026; Wang *et al*., 2023), and experimental protocols emphasising agronomic standards, and where the control is not only an untransformed wild type but also the best available commercial variety or hybrid (Sinclair and Rufty, 2022).

Factor analytical models could be insightful by linking with biological frameworks of phenotypic plasticity. The genotype scores from an FA model play the role of random regression slopes on latent environmental axes, and the environment loadings act as those axes. When factor-1 loadings align with environment quality, the first genotype score approximates the slope of a reaction norm. FA thereby decomposes genotype-dependent, phenotypic response to the environment into systematic and residual components. In a substantive sample of genotypes and environments where oat yield was measured in breeding plots, phenotypic plasticity was agronomically adaptive and correlated with overall performance calculated with FA (Fig. 3AC). In a similarly large, agronomically relevant data set of wheat yield, phenotypic plasticity was maladaptive and correlated negatively with overall performance calculated with FA (Fig. 3BD).

## Conclusion

Our biological-statistical synthesis shows that intricate statistical models, which are opaque except for practitioners, return phenotypic traits that contain information comparable to transparent reaction norms relating yield and environmental mean yield, thus connecting the biologically rich concept of phenotypic plasticity with advanced statistics.

### Contributions

VS, JT: conceptualisation, writing; VS: literature review, visualisation; MW: prospect theory; BS: statistical analysis; JH: oat data, statistical analysis; MR: wheat data; all authors provided input in writing the paper and all approved the ms.

#### Box 1.

Stability and balance in agriculture from the viewpoint of prospect theory

##### Reasons for an unexamined preference for balance and stability in agriculture

As early as the 1700s, mathematicians observed that people’s economic decisions did not align with the calculated values of expected outcomes but instead showed a willingness to sacrifice value to avoid uncertainty (Bernoulli, 1738, as cited in Seidl, 2013). von Neumann and Morgenstern (1953) operationalised this idea as the ‘certainty equivalent’, the amount of money a person would accept in lieu of a risky gamble. As a simple (if extreme) example, imagine a gameshow where a contestant has a 50/50 chance of winning $1M or nothing but the host offers a guaranteed $400K as an alternative to playing. Most people will prefer the certain option despite it being less valuable than the $500K expected value of the gamble. What an individual will choose depends on their own circumstances and how the problem is framed but this bias towards certain outcomes is widely seen in behavioural economics and is a core aspect of prospect theory (Kahneman and Tversky, 1979), which describes how people in general act when faced with risky choices – being risk averse and preferring lower value certain outcomes when faced with risky choices about potential gains but risk seeking when considering losses. The strength of these tendencies, however, is affected by cultural norms such as masculinity/femininity, individualist/collectivist, and uncertainty acceptance/avoidance - effects commonly considered in comparisons between countries (*e.g.*, Rieger *et al*., 2015) but which can also differ between groups within a country. For example, farmers often share a cultural milieu distinct from the broader population of a country and show a greater tendency towards collectivism due to their interdependence and reliance on cooperation (*e.g*., Uchida *et al*., 2019) and the levels of collectivism even differ between growers of particular commodities, *e.g.,* wheat or rice (Sadras and Hayman, 2025; Talhelm and Dong, 2024). Given this, the role of known decision-making tendencies and biases goes some way towards explaining why an unexamined preference for balance or stability might exist in agriculture.

##### A parallel between agronomically adaptive plasticity and ‘heuristics and biases’ in decision making

We advanced a definition of agronomically adaptive plasticity, which is best seen as a null-hypothesis (see *Agronomically adaptive plasticity*). This parallels Tversky and Kahneman’s (1974) ‘heuristics and biases’ approach to decision making, which remains central to behavioural economics and psychology. Here, the purely rational behaviour predicted by economic theories of behaviour (*e.g*., utility theory) is used as optimal standard and compared to actual real-world behaviour. The systematic differences between optimal and actual decisions are the ‘biases’ while the proposed cognitive explanations of why they occur are the ‘heuristics’. As an example, the bias commonly called base rate neglect is a tendency for people to underweight base rate information (*e.g.,* prior probabilities and long-term averages) when presented with both the base rate and new information (Tversky and Kahneman, 1974). Experiments observing conditions that strengthen and weaken this effect have identified constraints on human decision making based in how our memories and other cognitive processes work and, in particular, how these interact with environmental constraints such as non-stationarity, *i.e.,* variation in rates of occurrence across locations – to give rise to base rate neglect and limit how close to optimal human decision making can be (Bar-Hillel, 1980; Welsh and Navarro, 2012).

